# EMAP-SSN: An Embedding- and Multiple-Alignment-Integrated Sequence Similarity Network Platform for Interactive Exploration of Protein Sequence Space

**DOI:** 10.64898/2026.09.05.748609

**Authors:** Xuebin Feng, Emma R. Master

## Abstract

Sequence similarity networks (SSNs) are graphical representations of sequence relationship frequently used for exploring protein sequence space. Conventional SSN workflows typically use BLAST to calculate sequence similarities and rely on external visualization tools to generate the final networks. Consequently, raw sequence data is often detached from the calculated similarities during visualization, complicating SSN analyses that require residue-level information. To bridge this gap, we present EMAP-SSN, an open-source, cross-platform software suite that integrates SSN computation, visualization, and analyses in streamlined workflows. The program provides BLAST- and embedding-based alignment pipelines for sequence-similarity calculation and directly links network nodes to their original sequences and multiple alignments for analyses. Modular architectures for embedding generation, command integration, and browser-based utilities allow additions of research-specific functionalities and facilitate future development. Using a set of fold-type IV pyridoxal 5′-phosphate-dependent enzymes, we demonstrate how EMAP-SSN connects network topology with residue-level variation to identify sequence clusters, map functional motifs, and detect subgroup-specific conservation patterns. These capabilities provide a practical route from large protein sequence sets to experimentally verifiable hypotheses on enzyme function and targets for protein engineering. The EMAP-SSN program can be accessed from https://github.com/Xuebin-Feng/EMAP-SSN.

## Introduction

Sequence similarity network (SSN) is a bioinformatics tool designed to illustrate sequence-level similarities between proteins or genes using graph-based visualization [1], [2]. In an SSN, sequences are represented as nodes, and pairwise similarities above a user-defined threshold are shown as connecting edges. The final SSN layout is typically calculated via a force-directed algorithm, which simulates a physical spring-and-mass system to group sequence nodes into visual clusters [1]. Unlike other visualization tools such as uniform manifold approximation and projection (UMAP) and principal component analysis (PCA) which require dimensionality reduction and feature extraction to convert sequences to representative vectors [3], an SSN is based on raw sequence similarities computed via pairwise alignment, preserving high-dimensional complexity of pairwise sequence relationships in 2D clusters. Compared to phylogenetic trees, SSNs allow for the visualization of non-hierarchical connections among sequences and enable the visualization of thousands of nodes without overcrowding the canvas [1].

A typical SSN workflow consists of first calculating sequence similarities via all-vs-all pairwise alignment and then generating the SSN using visualization software such as Cytoscape [1], [2]. The all-vs-all similarities are often computed using the BLAST program as E-values, a statistical measure representing the expected number of matches with a given score occurring solely by chance within a dataset [4], [5]. As a heuristic algorithm optimized for rapid database searching to extract sequences sharing local similarities, BLAST applies asymmetric treatments to query and subject sequences [4]. This directional bias can lead to substantial differences in E-values depending on the choice of the query. The use of context-independent substitution matrices in BLAST causes the program to quickly lose accuracy when the sequence identity drops into the “twilight zone” (20% to 35%), missing potential matches among relatively distant clusters [6], [7]. On the other hand, the Karlin-Altschul statistics establish that the E-values are inherently dependent on dataset size; consequently, the E-value calculated for the same pair of sequences increases as the database expands [2], [8], creating problems for result reproducibility and scientific communication.

SSN layout generation in Cytoscape solely relies on node connectivity derived from sequence similarity with a user-defined threshold [1], [2]. Although relationships between sequences are being visualized, the underlying sequence information, such as amino acid frequencies and coevolution, is never passed to the visualization program. To bridge this gap, researchers have to obtain and format orthogonal information as "node attributes" and upload them for analysis, which requires considerable manual effort and data curation [2]. In some cases, researchers also need to have the original sequence set, or its multiple sequence alignment (MSA) on hand to check for locally conserved amino acids or to extract sequence features which are not immediately available in an SSN visualization [1]. This disconnection and loss of information between the computation and visualization steps complicates inline residue-level analyses and limits the biological interpretation of SSNs.

Recent developments in protein language models (pLMs) have enabled the accurate parameterization of protein sequences into numerical representations as embeddings while retaining abstract structural, functional, and biophysical property information [9], [10]. Per-residue protein embeddings are *L* × *D* matrices, where *L* is the sequence length (after stripping the special tokens) and *D* is the model-specific embedding dimension [11]. As the sequence length and amino acid order are preserved, embedding-based alignments (EBAs) have been explored as alternatives to traditional alignment algorithms, offering improved accuracy through context-aware representations, especially in the sequence identity “twilight zone” [7]. A recently published EBA algorithm by Pantolini et al. [12] employed exponential transform normalization and pseudo-Z-score signal enhancement to align embeddings by capturing contextualized residue-pair similarities. The method computes one similarity matrix for both Needleman-Wunsch (NW, “global”) and Smith-Waterman (SW, “local”) alignments using pure vector math and without a fixed substitution matrix, enabling potential parallelization on graphics processing units (GPUs).

To address limitations of conventional SSN pipelines, we present EMAP-SSN (Embedding- and Multiple-Alignment-integrated Protein Sequence Similarity Network), an interactive SSN computation, visualization and analysis platform developed to streamline SSN generation via both the BLAST- and EBA-based similarity calculation workflows while enabling residue-level inline analyses by integrating the original sequence set and its MSA. The program adapts and scales up the EBA algorithm by Pantolini et al. [12] (referred to as EBA hereafter) for GPU-accelerated sequence similarity calculations. An embedding-based MSA algorithm was developed based on the EBA to utilize pre-computed similarity networks for guide tree building, circumventing heuristic distance matrix calculations typical of conventional sequence-based methods. Driven by an extensible command-line interface (CLI) with direct MSA access, EMAP-SSN supports diverse inline analyses such as node selection, grouping, clustering and amino acid frequency profiling, outputting visual, textual and tabular results. The modular, plugin-style CLI architecture simplifies future development without the need to modify the core program framework. The EMAP-SSN program can help researchers to simplify and accelerate their protein discovery workflow and uncover new insights from existing sequence databases.

## Results

### Architecture Overview

EMAP-SSN is an open-source, Python-based application equipped with GPU-accelerated computational backends and interactive graphical user interfaces (GUIs) engineered for seamless cross-platform deployment on Windows, macOS, and Linux environments requiring minimal user setup. The EMAP-SSN source code is freely available on GitHub for academic and commercial use under Apache License 2.0.

Two typical EMAP-SSN workflows are illustrated in Figure 1A. The program is designed with two distinct but connected stages to isolate computational workloads from interactive visualizations and analyses. The computational stage is administered through the “Tools” GUI (Figure 1B), which manages protein embedding generation, all-vs-all sequence similarity calculation via EBA or BLAST, and embedding-based MSA building. The visualization and analysis stage is driven by the “Config” and “Viewer” GUIs (Figure 1C and 1D), which handle SSN layout calculations and serve as a workspace for interactive SSN visualization and analysis, respectively.

**Figure 1.**
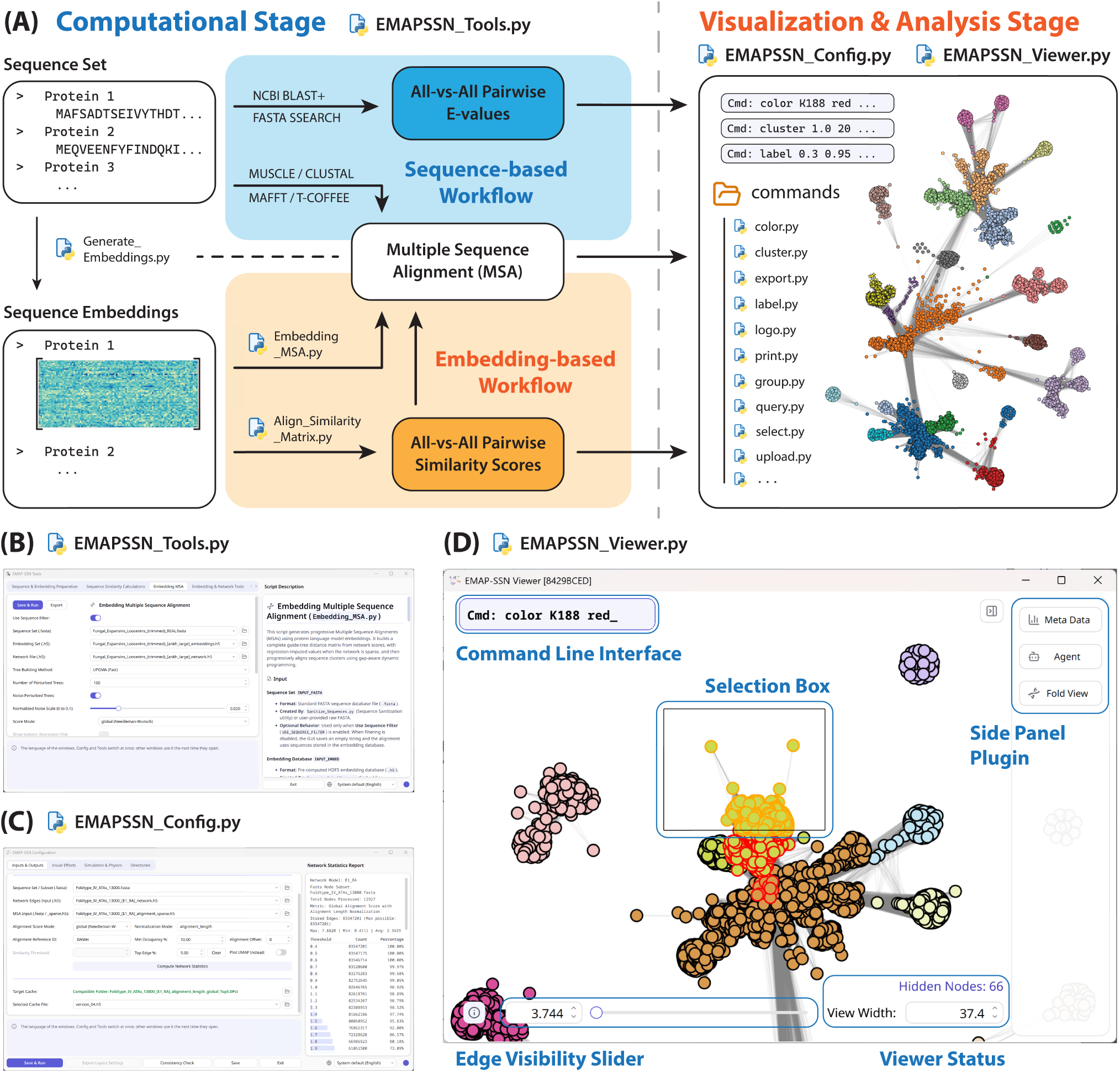
(A) Two typical EMAP-SSN workflows from a protein sequence set to the final SSN. (B) The “EMAP-SSN Tools” GUI handles the computational stage. (C) The “EMAP-SSN Config” GUI handles SSN plotting and cache loading. (D) The “EMAP-SSN Viewer” GUI handles the SSN visualization, interaction and analysis.

The computational stage features a dual-pipeline design. The “Tools” GUI handles an embedding-based workflow but also supports E-value calculations using a local BLAST executable. In an embedding-based workflow, each sequence in the sequence set is converted to an *L* × *D* per-residue embedding using the selected pLM model. The embeddings are pairwise aligned via EBA, and the similarity scores are calculated for each sequence pair. The embedding-based MSA uses both the sequence embeddings and the pre-computed similarities as inputs, performs progressive tree-based alignments, and writes the MSA in FASTA format. In a sequence-based workflow, the all-vs-all E-values are calculated using BLAST, and the MSA is obtained using an external MSA tool such as MUSCLE [13].

The visualization and analysis stage takes sequence similarity scores or BLAST E-values and an MSA as inputs. The “Config” GUI activates a layout engine that plots the SSN by simulating a spring-and-mass system. When a dynamic equilibrium is reached, individual components are packed into a Boolean grid with larger components at the center. The final SSN is rendered by VisPy and displayed in the “Viewer” GUI, which supports interactive component selection and manipulation, a CLI overlay supporting advanced formatting and analysis, and a side panel with registered links to browser-based utilities hosted by a local HTTP server.

### Capabilities and Example Analyses

To demonstrate the capabilities of EMAP-SSN, here we present an example workflow for exploring the sequence space of fold-type IV pyridoxal 5′-phosphate (PLP)-dependent enzymes. This functionally diverse fold-type comprises distinct, characterized enzyme groups, which include R-selective amine transaminases (R-ATAs), D-amino acid aminotransferases (DAATs), branched-chain amino acid aminotransferases (BCATs), and 4-amino-4-deoxychorismate lyases (ADCLs), which are distinguished by characteristic active-site signatures associated with respective catalytic functions and substrate recognition mechanisms [14], [15].

#### Example Dataset

The majority of the sequences were obtained from the omega-transaminase engineering database (oTAED). Despite having “omega-transaminase” in its name, the dataset contains enzymes with other functions under the same fold type. Additional sequences from the PDB database were added as structural references for analyses [16], [17], and the final sequence set consists of 12927 sequences with an average length of 297.5 amino acids.

#### Computation Stage

To ensure compatibility with downstream processes, the sequence set was pre-processed to substitute illegal characters in sequence headers and unsupported amino acid symbols in the sequences. The per-residue protein embeddings were generated using the ESMC 6B [18] model and the Profluent-E1 600M model in retrieval-augmented (RA) mode [19]. The global and local alignments were performed following the EBA, and the alignment/similarity scores were recorded. Multiple alignments of the sequence set were constructed using the embedding-based MSA, in which pairwise similarity scores obtained from the EBA were used to build guide trees. We also quantified sequence similarities using BLAST as E-values and aligned the sequence set using MUSCLE in super5 mode, which are used as analysis baselines.

#### Network Generation and Interaction

The SSNs were generated using the top 5% normalized sequence similarities following force-directed layout calculations. For EBA-derived similarities, alignment-length-normalized global alignment scores were used. For BLAST, no normalization was applied, and the cutoff corresponded to an E-value of 3.8 × 10^−99^. Node groups with no connecting edges are calculated independently and packed into square grids using macro-grid Boolean packing. As EMAP-SSN supports interactive layout adjustment, we manually repositioned node groups to reduce over-crowding or large blank areas on the canvas (**Figure 2**).

**Figure 2.**
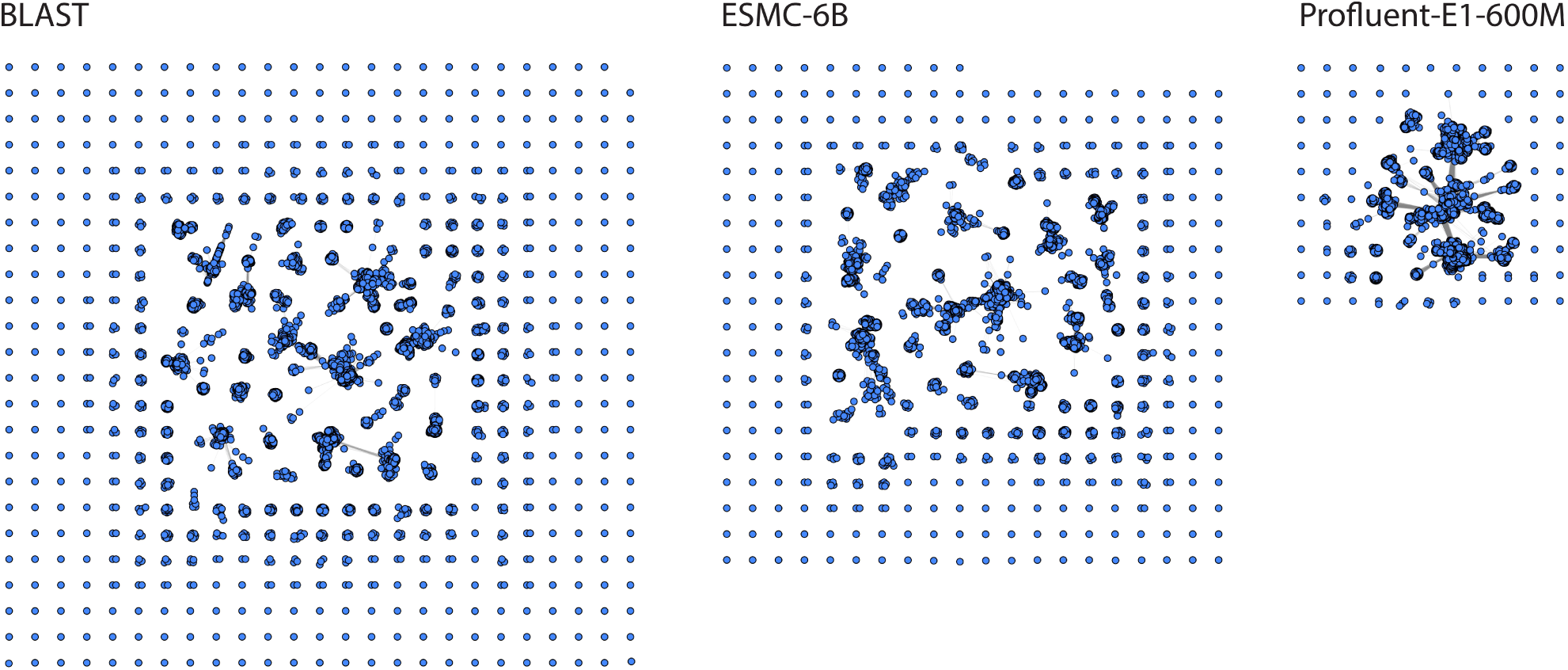
A comparison of SSN layouts generated using BLAST E-value and normalized EBA similarity scores calculated from embeddings derived from different pLM models.

Compared to the conventional BLAST-based SSN, EBA-based SSNs showed substantially greater connectivity among visual node clusters, supporting the previously reported higher sensitivity toward remote sequence relationships in the sequence-identity “twilight zone”[12]. The BLAST-based SSN contained the largest number of isolated nodes and small connected components, indicating that many pairwise relationships among node groups were not detected at the selected threshold. The ESMC-based SSN showed a broadly similar topology but contained larger connected components and substantially fewer free nodes. In contrast, the E1-based SSN was dominated by a single connected component containing more than 95% of the sequences, revealing putative relationships among sequence subgroups that remained disconnected in the BLAST- and ESMC-based networks. The increased connectivity may partly result from E1’s retrieval-augmented architecture, which can capture richer structural information by explicitly incorporating homolog sequence information into embedding representations. The E1-based SSN was used for the subsequent analyses.

#### Analyses via CLI Commands

EMAP-SSN provides a modular, plugin-style CLI overlay on the “Viewer” GUI for interactive, in-line analyses. Although the E1-based SSN layout showed distinct locally dense regions, cluster membership remained ambiguous for nodes in between or at the periphery of visual clusters. We thus applied the CLUSTER command to the E1-based SSN to use Leiden community-detection algorithm [20]. The detection was performed on the weighted network graph, using edge connectivity and associated sequence-similarity scores, and was independent of the distances between nodes in the visual layout. Using modularity optimization with a resolution of 1.0 and a minimum cluster size of 20 nodes, the command identified 21 clusters comprising 12,827 of the 12,927 nodes (99.23%) (**Figure 3A**). The remaining 100 nodes (0.77%) were categorized as noise.

**Figure 3.**
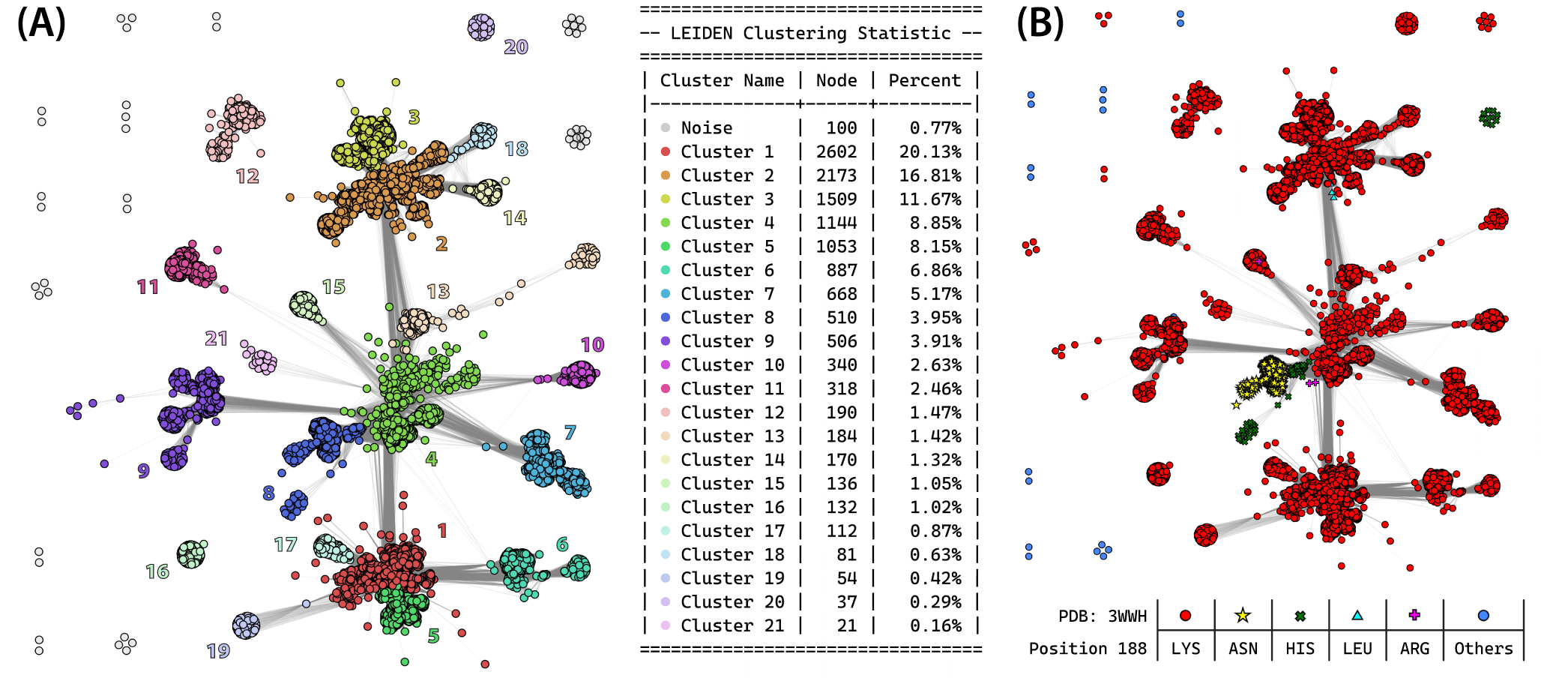
(A) The E1-based SSN (with free nodes stripped) colored by clusters using the CLUSTER command. The modified terminal table output is also shown. (B) The E1-based SSN with different colors and shapes assigned to nodes based on their conserved position amino acids.

Most characterized PLP-dependent enzymes contain a conserved active-site lysine that forms a covalent Schiff base with the 4′-aldehyde of PLP, thereby helping to retain the cofactor in the resting enzyme. In transaminases, this lysine additionally functions as a general acid-base catalyst during proton transfer steps and is therefore important for efficient catalysis. The previous study by Buß et al. [16] reported that fold-type I enzymes have this position conserved to nearly 100% lysine. In fold-type IV, however, only ∼ 88% of sequences are reported to have lysine at the corresponding conserved position, raising questions of catalytic competence and physiological functions of fold-type IV proteins lacking this canonical residue.

Since EMAP-SSN associates each node in the SSN with its corresponding sequence in the MSA, we were able to quickly identify nodes that are missing the conserved lysine and query what they have at the equivalent position using a list of CLI commands. First, we selected the R-ATA from *Arthrobacter sp.* KNK168 (PDB: 3WWH) as the MSA reference using the REFERENCE command, thereby expressing alignment indices according to the residue numbering of a known sequence. We then selected nodes that do not have lysine at the conserved position by passing the Boolean expression !K188 to SELECT. The QUERY command was subsequently used to determine the residue frequencies at the conserved position of selected sequences, and different colors and shapes were assigned to all sequences based on their conserved position amino acid using COLOR, generating the SSN shown in **Figure 3B**.

Using the embedding-based MSA calculated from the E1 embeddings, analysis showed that ∼ 95% of sequences contain the conserved lysine, much higher than the ∼ 88% reported by the previous study [16]. Sequences lacking the conserved lysine have mainly asparagine (∼ 80.8%) and histidine (∼ 13.1%) and are likely closely related as they are mostly assigned to cluster 8. To perform the same analysis with the MSA constructed using MUSCLE in super5 mode, we used the ALIGNMENT command to load the MSA without replotting the SSN, which showed nearly identical results (Figure S1).

Beyond the alignment-based analyses, the CLI also supports metadata-based queries and modifications to the SSN visualization. During the program initialization, the “Viewer” GUI registers its bundled web utilities and launches a local, lightweight HTTP server. Calling the META command opens a Tabulator-based viewer in user’s default browser, from which metadata can be imported from Excel or CSV files into the active session by exact header matching. For a newly calculated layout, the EMAP-SSN program derives sequence lengths from the configured FASTA records and creates “Length” as a default metadata column. Using the SPECTRUM command to color nodes by their sequence length, we found that a subset of plant-sourced sequences that have histidine replacing the conserved lysine are substantially longer than rest of fold-type IV sequences on average (524.5 vs 297.5 aa) (**Figure 4**). We then used the ESMFOLD command to predict the structure of such a sequence (supplementary material) using an API-hosted ESM3 (98B) model, and structurally superposed the predicted structure onto 3WWH in the integrated Mol* web interface [21]. The predicted structure consists of an extended N-terminal segment connected via a flexible linker to a C-terminal region comprising the canonical fold-type IV two-domain architecture. The additional N-terminal extension was annotated by InterPro as a sulfotransferase 5 domain (PF19798) [22]. A BCAT from tomato plant (UniProt: A0A3Q7F7J3) has been reported to have a similar architecture and exhibit canonical BCAT activities [23]. However, like other cluster 8 sequences, the enzyme lacks the conserved catalytic lysine, leaving its mechanistic basis of reported activity unclear. Five proteins from *Burkholderia* sp. in cluster 9 similarly contained long N-terminal extensions, which are annotated by InterPro as peptide deformylase domains (PF01327) (Figure S2A). These extensions retain key residues of the conserved catalytic center of peptide deformylases, suggesting that they might be catalytically competent rather than just nonfunctional additions (Figure S2B) [24].

**Figure 4.**
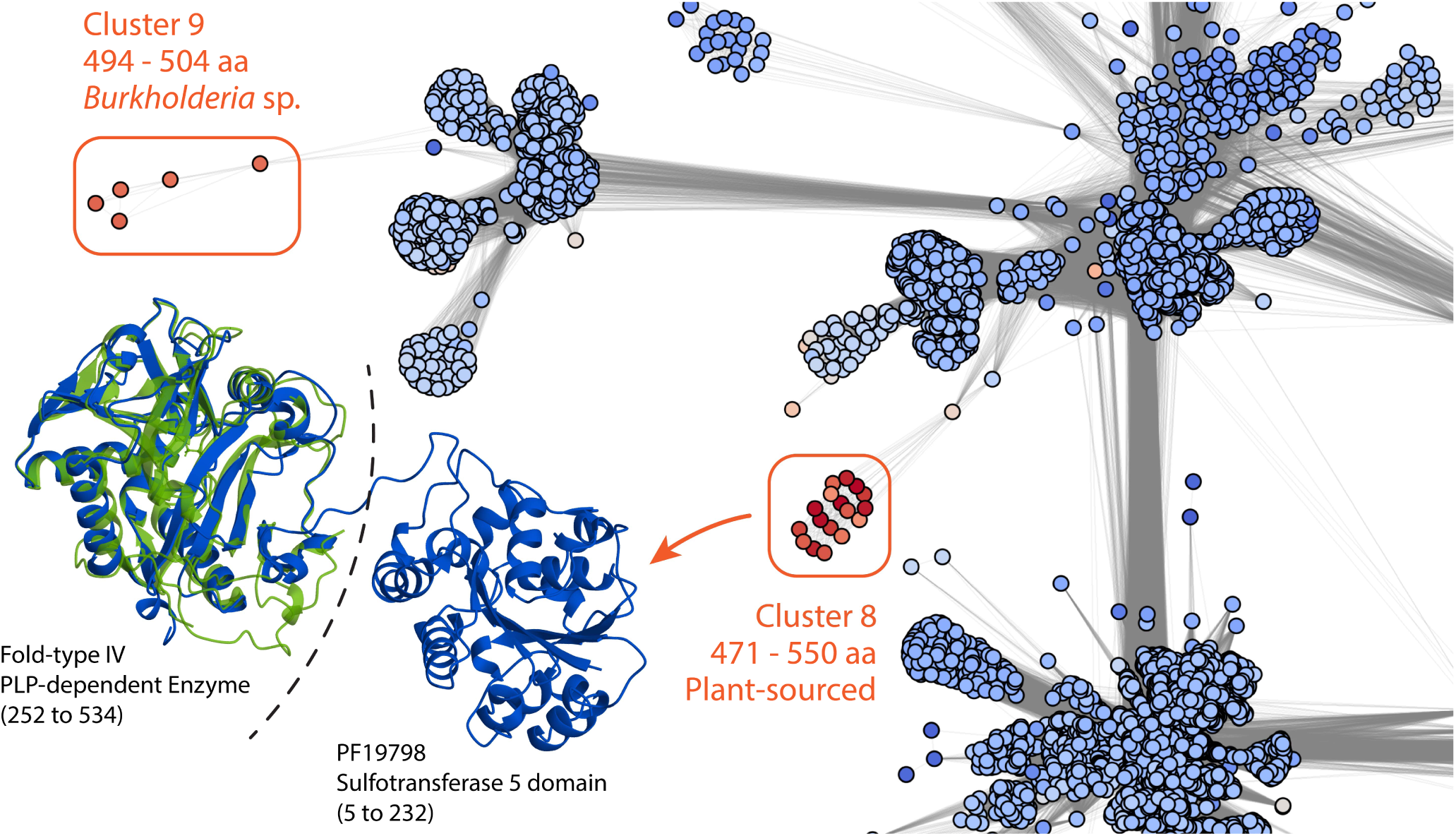
The E1-based SSN colored by sequence length metadata using the SPECTRUM command. Two groups of unusually long sequences are highlighted. The structural alignment of *Glycine soja* protein KHN16146 against the 3WWH reference is shown in the inset. The predicted KHN16146 structure is colored based on pLDDT and the 3WWH structure is colored in green.

Previous studies by Höhne et al. [15] identified functional-group-associated active-site residue signatures of fold-type IV PLP-dependent enzymes by analyzing their structural characteristics and substrate-recognition mechanisms (**Figure 5A**). Although useful in classifying and predicting functions of uncharacterized sequences, the short active-site motifs do not capture broader sequence relationships among functional groups, or group-specific residue conservation outside the active site. Using EMAP-SSN, we mapped reported residue signatures to the E1-based SSN, enabling visualization of functional-group relations, investigation of amino acid conservation, and identification of uncharacterized sequences connected to proteins with established functional groups despite lacking key active-site residues. By sequentially executing the commands shown in **Figure 5A** in the CLI, the SSN shown in **Figure 5B** was obtained. Note that canonical DAAT (C-DAAT) and non-canonical DAAT (NC-DAAT) have different active-site signatures, which were summarized by Shilova et al. [25]. **Figure 5B** was generated using the ESMC-derived MSA on the E1-based SSN, since the E1-derived MSA did not align 3WWH positions 135 and 138 to corresponding active-site signature of C-DAAT. Our analyses show that this discrepancy is unlikely a problem of the embedding-based algorithm; rather, it shows that these positions in the reference sequence 3WWH cannot be reliably aligned to those in C-DAATs (Figure S3).

**Figure 5.**
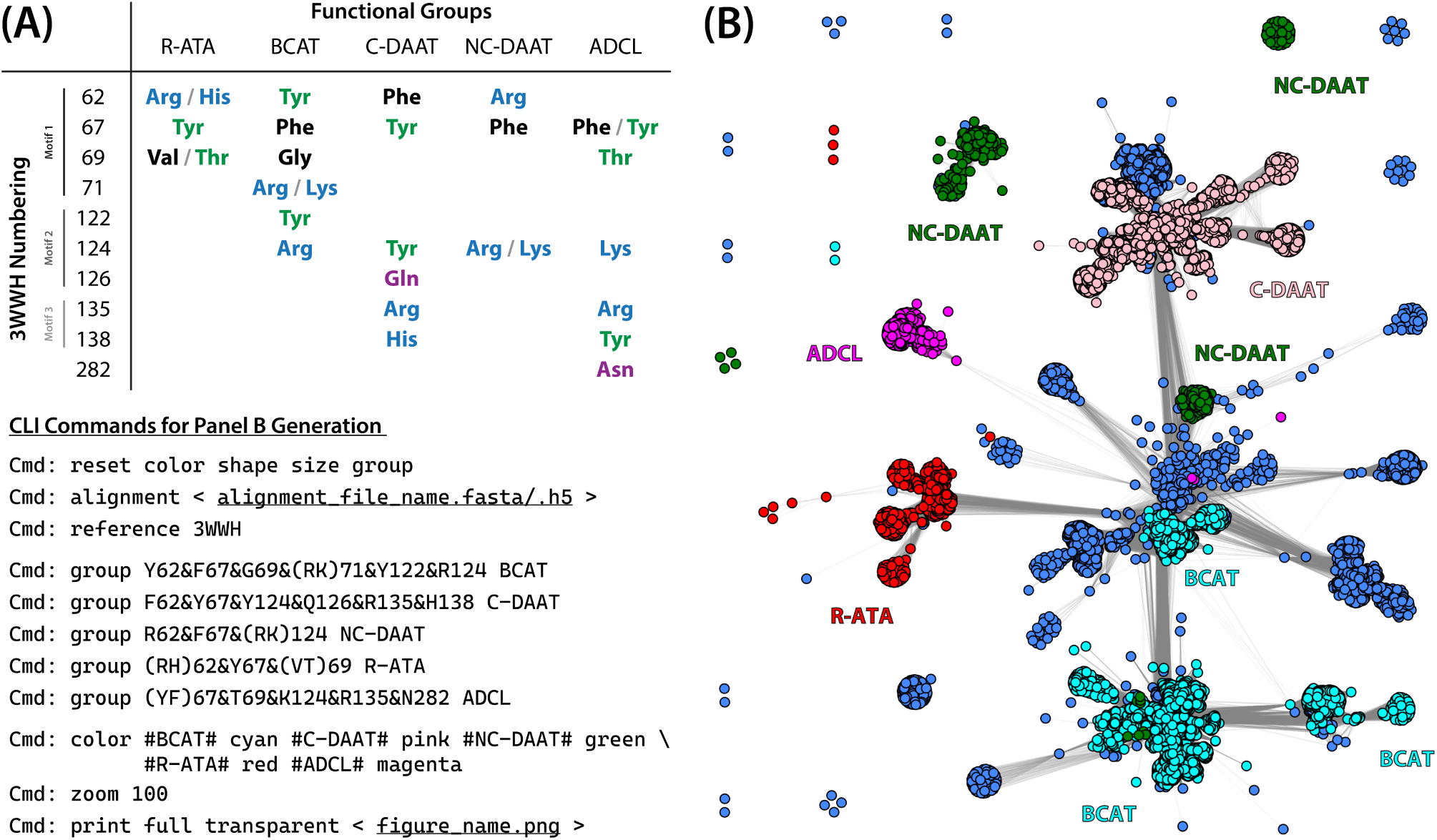
(A) Function-group-associated residue signature identified by Höhne et al. [15] adapted to 3WWH numbering scheme with corresponding CLI commands used to generate panel B. (B) E1-based SSN colored based on active-site signatures of sequences.

Interestingly, the signature-defined functional groups occupied distinct peripheral regions in the E1-based SSN but remained connected to central clusters that consist mainly of sequences with motifs matching none of the active-site signatures. As expected, sequences in cluster 8, which lack the conserved lysine, do not have signature residues of any characterized functional groups. Whereas R-ATAs (cluster 9) and ADCLs (cluster 11) separately form dense and self-contained clusters with well-defined boundaries, sequences matching DAAT and BCAT signatures are distributed across multiple clusters, or are in close association with sequences that do not share the same active-site motifs.

To investigate what additional residues are conserved within each functional group beyond the defining signatures, we used the LABEL command to detect subset-specific conservation patterns that are not globally significant. The command reads the active MSA and current cluster and group settings, filters the MSA to construct subset alignments, and compares local (per subset) and global amino acid frequencies to generate a formatted XLSX output. Here, we studied the five functional groups and two custom groups forming cluster 8 that have asparagine (Active-ASN) or histidine (Active-HIS) at the conserved lysine position. The command was used to highlight locally conserved (≥97% in a group) residues that have below 20% frequencies after excluding groups with the same local conservation from the global context.

The analysis revealed additional group-specific conserved positions featuring highly divergent residue profiles from the global background. Using the LOGO command, we plotted the sequence logos of each group at these positions in weighted bits to visually illustrate their local conservation (**Figure 6A**). Outside the three signature motif regions, R-ATA conserves N189 and D194 on the N-terminal side of the catalytic K188 in the PLP binding pocket; C-DAAT and ADCL both conserve R58 near the homodimer interface, whereas ADCL additionally conserves G242 at the C-terminal end of a flexible loop. All five functional groups are strongly conserved at position 192, which corresponds to a cofactor binding site residue pointing to the 3′-hydroxyl of PLP (Figure S4A). Interestingly, this conservation is group-specific, with five functional groups conserved to four different amino acids. Using the QUERY and COLOR commands to color the E1-based SSN according to the amino acid at this position, we found that the local residue identity is strongly associated with previously identified node clusters (Figure S4B). Active-ASN and Active-HIS, which lack the conserved lysine, strongly conserve rare amino acids W67 and H122 near the cofactor- and substrate-binding domains. These conservations may offer clues to the cofactor-binding modes and natural functions of these proteins, although whether they covalently bind the PLP remains unexplored. The two groups also share a rare but highly conserved N223 at the homodimer interface; the Active-ASN additionally conserves P135 and Q138 in motif 3 while the Active-HIS conserves H189 right beside its defining residue H188; the functions of all three residues are unclear.

**Figure 6.**
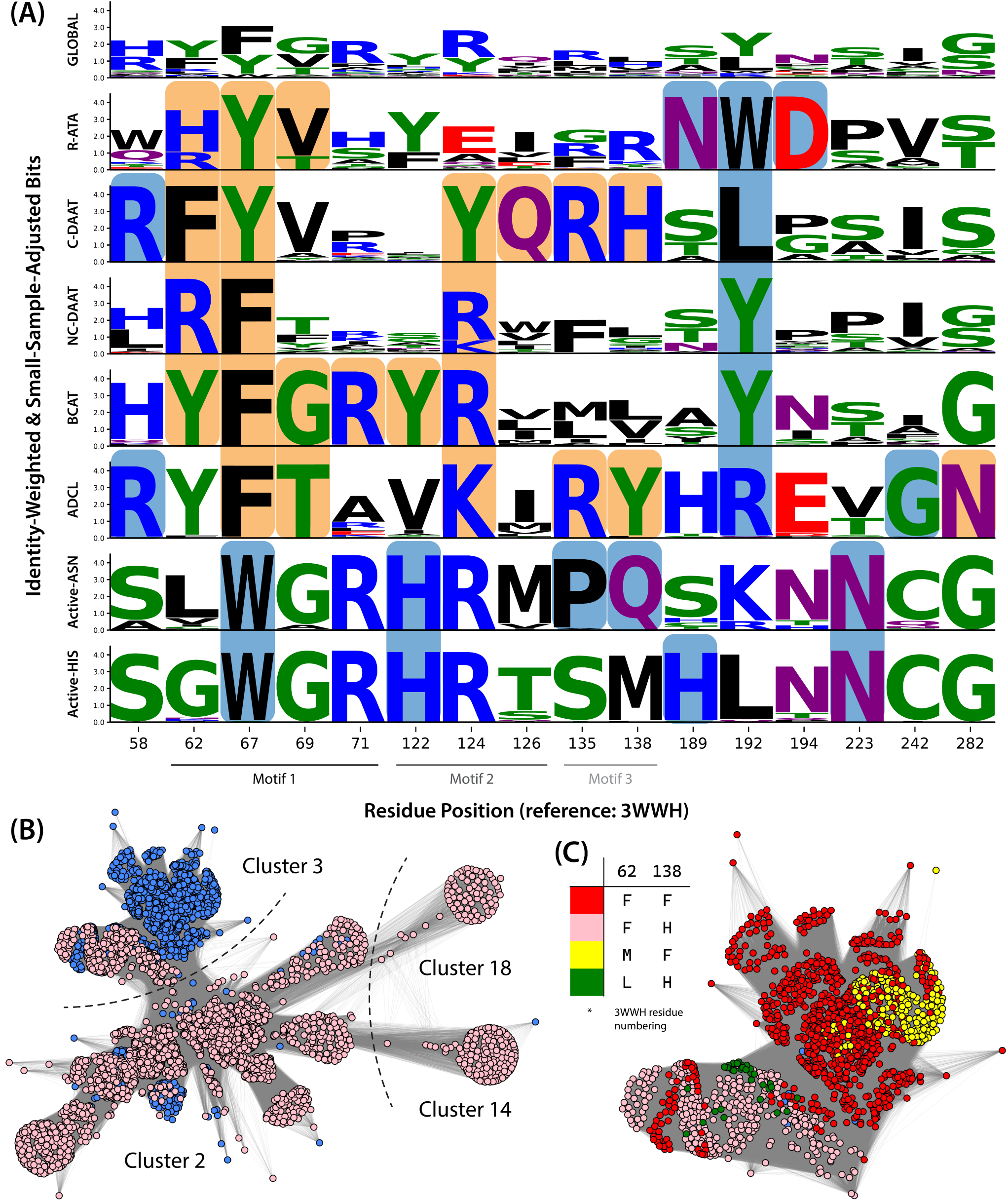
(A) Sequence logos generated using the LOGO command at signature positions and additional positions of interest identified using LABEL, highlighted with orange and blue backgrounds, respectively. Residue positions are numbered using 3WWH as the reference. Information content (bits) was corrected for small sample size and weighted by sequence identity. (B) The four clusters containing C-DAAT. Nodes colored in pink match the C-DAAT active-site signature. (C) Cluster 3 nodes colored based on the amino acids at positions 62 and 138. Not all nodes contain combinations of amino acids illustrated in the table.

Clustering of C-DAATs showed a distinct topology. Whereas clusters 2, 14 and 18 contain mainly predicted C-DAATs, cluster 3 includes a large portion of sequences which do not match the CDAAT active-site motif (**Figure 6B**). Checking signature positions using QUERY, we found that Y67, Y124, and R135 were strongly conserved among cluster 3 sequences, consistent with the C-DAAT motif. Glutamine predominated position 126, with glutamate also observed (Q: 94.4%; E: 4.2%). Greater variations occur at positions 62 (F: 84.3%; M: 13.4%; L: 2.3%) and 138 (F: 76.1%; H: 23.6%), for which we used the COLOR command to visually illustrate their amino acid distributions (**Figure 6C**). In C-DAAT, F62 is mainly a hydrophobic, pocket-shaping residue in the first coordination sphere of the active site instead of a direct coordinator of the substrate α-carboxylate [26]. A substitution by methionine or leucine will retain side-chain length but likely alter substrate-binding pattern due to the missing aromaticity. The H138 in C-DAAT is part of the “carboxylate trap” critical for substrate recognition [27]. A substitution to phenylalanine will preserve the side-chain length, aromaticity, but not polarity. This might alter substrate scope and induce promiscuous R-ATA activity due to weaker specificity toward amines with α-carboxylate groups. Further, we noticed that M62 does not coexist with H138 in the sequence set. As both positions are within the C-DAAT substrate-binding environment, the absence of the M62/H138 combination may reflect a constraint on compatible amino acid pairs. Although M62 and H138 each occur in cluster 3, their combination may be unfavorable for substrate binding; however, this hypothesis requires more detailed covariance analysis or experimental validation. Such sequence-level patterns identified through the EMAP-SSN program can help prioritize divergent enzymes for experimental characterization and guide selection of residues for protein engineering. Lists of commands used to generate all SSN and sequence logo figures are available in the supplementary material.

### Embedding-based MSA Benchmarking

In the above example, accuracy of MSA is an important determinant for the validity of analyses that rely on residue-level, positional homology. Whereas all three alignments correctly aligned the conserved active-site lysine, E1- and ESMC-derived MSAs disagreed on the alignment of the C-DAAT active-site motif, which can lead to errors in the analyses results.

To compare the embedding-based MSA against conventional sequence-based methods, we additionally constructed MSAs of the same sequence set using CLUSTALO [28], MAFFT [29] and embedding-based algorithm with embeddings generated using Ankh (base (450M) and large (1.2B)), ProstT5 (encoder-only, 1.2B), ProtBERT (420M) and smaller ESMC (300M and 600M) models. The MSAs were compared based on the sum of pair F1 (SP F1) [30] and the total column (TC) [31] scores against a reference set of 27 sequences aligned using the Expresso structural alignment in the T-COFFEE suite [32]. The two alignment scores, positions of total-column recovery, as well as the alignment times are shown in **Figure 7**.

**Figure 7.**
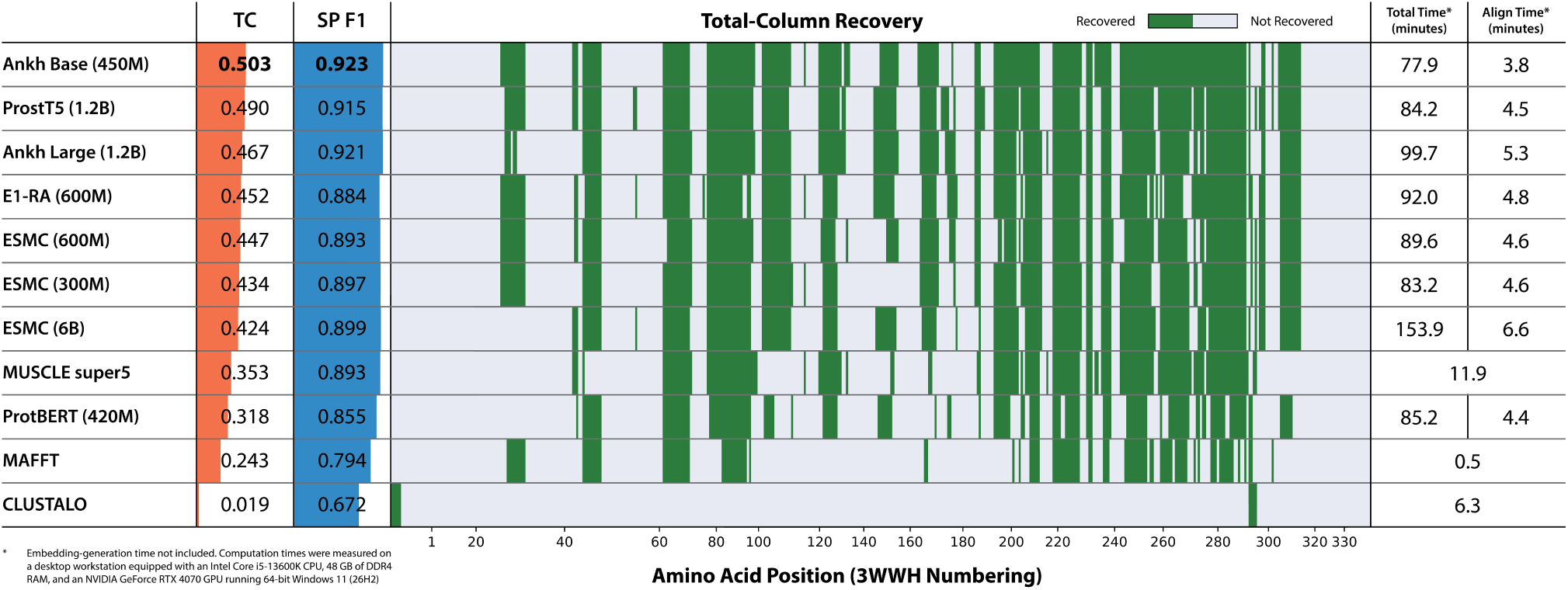
Total column (TC) and sum of pair F1 (SP F1) scores of fold-type IV PLP-dependent enzyme MSAs calculated using embedding-based and traditional algorithms. The positions of total-column recovery are shown as a heatmap with 3WWH residue numbering. The total and alignment-only times of MSA construction were measured with and without counting for the time spent on all-vs-all sequence similarity score calculations, respectively.

The embedding-based alignments generally outperformed those generated with conventional methods on both TC and SP F1 scoring metrics with MUSCLE super5 MSA ranked only slightly above the embedding-based MSA calculated using ProtBERT embeddings. Unexpectedly, the Ankh base (450M)-derived MSA achieved the highest scores in both metrics despite its small parameter size. A larger pLM model is not guaranteed to produce a more accurate MSA using the embedding-based MSA algorithm. When comparing computational times, however, the sequence-based algorithms showed a clear advantage as the embedding-based MSA requires pre-computed all-vs-all sequence similarities to build the guide trees. The high computational cost is justifiable if the embedding-based MSA is included as a part of the SSN workflow. For a sequence-based SSN workflow, MUSCLE super5 provided the strongest accuracy–runtime balance among the conventional methods tested.

## Discussion

### Improvements over the Conventional SSN Workflow

The EMAP-SSN program provides EBA as an alternative to BLAST for sequence similarity calculations and directly links each node to its corresponding sequence in an MSA, addressing the separation between similarity calculation, network visualization, and residue-level analysis in conventional SSN workflows. In the fold-type IV PLP-dependent enzyme example, embedding-based SSNs showed a more interconnected topology, demonstrating the advantage of EBA in capturing remote sequence relationships. Calling a sequence of commands through the CLI in the “Viewer” GUI, we interactively divided the SSN into clusters based on pairwise similarities, examined the conserved lysine distribution, identified functional groups based on characterized active-site signatures, and detected additional group-specific conservations from the MSA, which could be missed in structural or mechanistic studies. By combining direct network interaction with MSA-based selection, grouping, visualization, and analysis, the EMAP-SSN program enables relationships between network topology and residue-level variation to be investigated in one environment. This integrated workflow can facilitate hypothesis generation and the prioritization of sequences or residues for subsequent structural, mechanistic, and experimental studies.

### Scalability and Accuracy Limitations

The principal scalability limitation of the embedding-based workflow is the cost of all-vs-all residue-level EBA, as demonstrated in Figure 7. For a set of *N* sequences, a complete, undirected similarity network requires *N*(*N* − 1)/2 pairwise alignments. For each alignment, the similarity matrix scales linearly with model-specific embedding dimension *D* and quadratically with the average sequence length 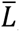, bringing the combined computational complexity to 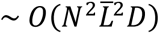. To mitigate this problem, EMAP-SSN supports the generation of sparse similarity networks consisting of only high-similarity pairs ranked by length-normalized cosine similarities of pooled embeddings. The sparse networks generally produce SSNs with similar topologies to that of the full network (Figure S5) for the fold-type IV enzyme sequence set; however, this approximation can negatively impact embedding-based MSA calculation (Figure S6), as the latter relies on isotonic regression between the cosine similarities and real pairwise alignment scores to impute missing similarity values (Figure S7).

As demonstrated in the example, many analysis commands in EMAP-SSN require an accurate MSA to prevent invalid results. However, MSA algorithms can rarely reproduce all columns present in a curated reference, as shown in Figure 7, and a set of divergent sequences may not be amenable to a single reliable MSA. Thus, we strongly recommend that users assess the robustness of their analyses by aligning representative protein structures and verifying that residues of interest occupy structurally corresponding positions. When experimental structures are unavailable, the ESMFOLD command can be used for structure prediction with ESM3 and load the resulting backbone models into the integrated Mol* web interface for structural superposition. These predictions should be interpreted in light of their confidence (pLDDT) and used as supporting rather than definitive evidence.

### Extensibility and Future Developments

The EMAP-SSN platform was developed with extensibility as a guiding principle. Its embedding-generation pipeline, CLI and browser-based utility modules were built with plugin-style architectures, allowing support for new pLMs, commands, and web-based utilities to be added without extensively modifying the codebase. For an embedding-based workflow, quality of the pLM model fundamentally affects the SSN topology and embedding-based MSA accuracy. With an upgradeable pipeline, sequences can be represented using increasingly more accurate and biologically informative representations as new pLMs are developed while maintaining the existing workflows. The modular CLI allows research-specific functionality to be implemented as independent command modules. A centralized dispatcher loads the modules dynamically and passes each command the active “Viewer” GUI instance, providing access to the network state, MSA, metadata, and visualization controls, while a shared command engine supplies common parsing and analysis functions. The “Viewer” GUI also hosts a local web server that automatically discovers validated descriptors for bundled web utilities and registers their routes, actions, and state contributions at startup. Existing utilities include a Tabulator-based metadata viewer and editor (Figure S8A) for displaying and modifying imported metadata; a Mol* web interface (Figure S8B) for visualizing structures loaded locally or predicted using the ESMFOLD command; and an agent chat panel (Figure S8C) which connects to a local or remote API and translates natural-language requests into corresponding CLI commands. By facilitating the integration of emerging bioinformatics tools and methods, this extensible architecture will help the program remain adaptable and relevant as the field continues to evolve.

## Methods

### Program Design

EMAP-SSN is a Python-based program built on open-source packages and is released on GitHub under the permissive Apache License 2.0. The program is compatible with Windows, macOS and Linux operating systems and executes in an isolated Python 3.12 environment containing required dependencies, which are installed automatically at program start. The three main GUIs were built with PySide6 (Qt 6.9) with the nodes and edges of the SSN rendered with VisPy. The JavaScript-based web GUIs are delivered through a lightweight local HTTP server, opened in the system’s default browser, and synchronized with the “Viewer” GUI to support interactive data exchange. The computational backend of EMAP-SSN provides separate CPU and GPU execution paths, using Numba just-in-time compilation for CPU kernels and PyTorch for certain operations that can run on either CPU or GPU. The optimal device is, by default, automatically selected based on user hardware via benchmarking prior to heavy computations to ensure optimal efficiency.

### Sequence Set Sanitization

The sequence set sanitization is handled by *Sanitize_Sequences.py* and is necessary to ensure compatibility with pLM models and CLI commands. FASTA headers are standardized by replacing characters that may interfere with command-line parsing or file handling, while amino acid sequences are converted to uppercase, terminal non-residue characters are removed, and unsupported internal characters are replaced with “X”. Empty records and duplicate sequences are removed, conflicting headers are resolved, and records can optionally be filtered according to user-defined sequence-length limits or header text.

### Per-residue Embedding Generation

The per-residue protein embedding generation is handled by *Generate_Embeddings.py* via the “Tools” GUI. The script was designed with a plugin-style pLM model library to improve flexibility and simplify future upgrades. Natively supported models include ESM2, ESMC, Ankh, ProstT5, and ProtBERT [9], [18], [33], [34], [35], with embeddings computed locally or through an API. The embedding generation script uses sequence sets in FASTA format as inputs, passes individual sequences to the pLM backends and receives per-residue embeddings in 32-bit floating-point precision with special tokens stripped. The script then converts the embeddings to the user-selected numerical precision and compiles the result in HDF5 format.

### EBA and Similarity Score Calculations

The EBA algorithm handled by *Align_Similarity_Matrix.py* is adapted from the work of Pantolini et al. [12] and implemented using a cosine-similarity-based metric with user-configurable gap penalties. Briefly, the similarity matrix of two per-residue embeddings calculated from sequence *A* and *B* of lengths *M* and *N*, respectively, can be calculated as:

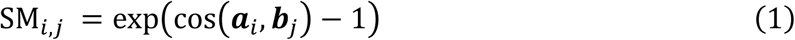

where ***a****_i_* and ***b****_j_* denote the embedding vectors of the *i*-th residue of sequence *A* and the *jj*-th residue of sequence *B*, respectively, and SM*_i_*_,*jj*_denotes the element in row *i* and column *jj* of the similarity matrix SM.

The row and column Z-score-standardized similarity matrices can be calculated as:

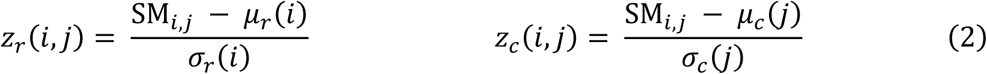

where the *μ_r_*(*i*) and *μ_c_*(*j*) are row and column sample means defined as:

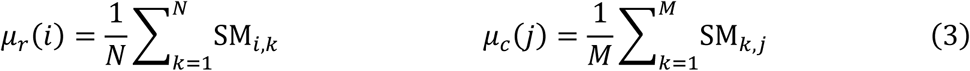

and the *σ_r_*(*i*) and *σ_c_*(*j*) are population standard deviations defined as:

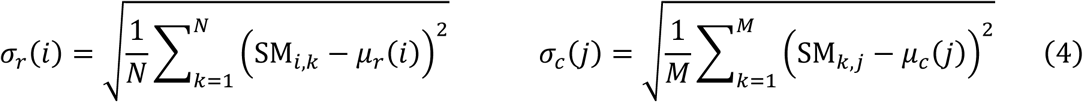

The final enhanced similarity matrix is calculated as:

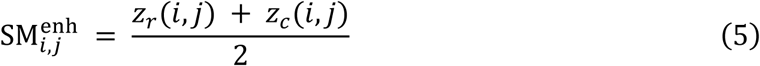

Global and local alignment scores are calculated concurrently using the standard NW and SW dynamic programming recurrences with user-configurable gap penalties (default values of 0 and −2.0 for global and local alignment, respectively). For local alignments, a constant value of 2 is subtracted from each element of the enhanced similarity matrix before applying the SW recurrence. The score and length of the optimal path were propagated together using rolling dynamic programming rows without constructing the full traceback paths. The resulting global and local alignment scores, corresponding alignment lengths, and sequence-pair indices are stored in HDF5 format.

For an all-vs-all network, EBA scores were calculated for all *N*(*N* − 1)/2 unique sequence pairs. An optional prefilter is provided to reduce the number of full residue-level alignments for large sequence sets. Each sequence embedding is first reduced to a single vector by max (default) or mean pooling, and the estimated similarity between sequences *A* and *B* can be calculated as:

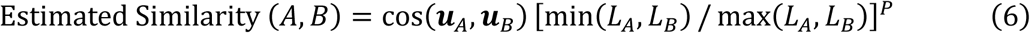

where ***u****_A_* and ***u****_B_* are the pooled sequence embeddings using the selected method, *L_A_* and *L_B_* are sequence lengths, and *P* (default 2) is the user-defined length-ratio exponent. Sequence pairs within the lowest user-defined percentile of adjusted similarities are excluded, and full residue-level EBA scores are calculated and stored only for the remaining pairs.

### SSN Layout Generation

The SSN layouts are generated from edges with normalized similarity scores (or negative log-transformed E-values) above a user-selected threshold using a spring-and-mass force-directed layout based on configurable physics parameters. Nodes are modeled as unit-mass particles with velocities whereas edges are represented as attractive zero-rest-length springs. Repulsive interactions and damped Euler integration separate the nodes and relax the layout. Each connected component is treated as an independent object. Components containing at least 500 nodes are simulated individually. Components containing fewer than 500 nodes are combined into computational batches with individual components remain isolated from one another. A batch can contain at most 2,000 nodes. For sufficiently large components having more than 2000 nodes, optional edge-based staged annealing can be applied to relax stronger connections first and then progressively introduce weaker connections until the similarity threshold is reached. After the simulation, nodes and edges of each component are converted into a Boolean occupancy mask, and component masks are packed without overlap into a global grid to produce the final SSN layout.

### Embedding-based MSA from Pre-computed Similarity Scores

Embedding-based MSA construction is handled by *Embedding_MSA.py* via progressive profile alignment guided by pre-computed normalized similarity scores (or negative log-transformed E-values). Similarities are first converted to distances by:

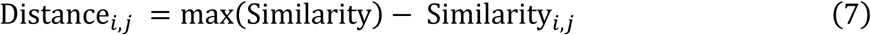

where *i* and *j* denote the indices of the sequences compared. This sets the lower bound of distance to 0, and sequence pairs with higher similarities are assigned shorter distances. A guide tree is then built from the calculated pairwise distances using either UPGMA or the neighbor-joining algorithm, with an optional consensus procedure based on noise-perturbed replicates. When enabled, zero-mean Gaussian noise is added independently to the distance matrix for each replicate, and the final consensus guide tree is constructed from the averaged cophenetic distances of the replicate trees.

Progressive alignment is then performed by traversing the final guide tree from leaves to the root. Before alignment, each residue embedding is L2-normalized. A profile column is represented by the sequence-count-weighted mean of the normalized residue embeddings at that position, with gaps contributing zero vectors. Consequently, the magnitude of a profile vector reflected both column occupancy and agreement among its residue embeddings. For profiles *A* and *B*, the directional cosine similarities between profile columns were transformed using the same combined row- and column-wise Z-score standardization used for the EBA similarity matrix (Equations 1 to 5). The final profile score was calculated as:

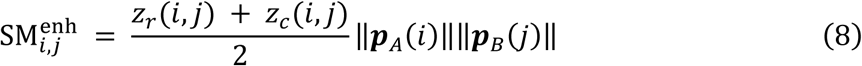

where ***p****_A_*(*i*) and ***p****_B_*(*j*) are the vectors representing columns *i* and *j*, respectively. The profile-norm term reduced the contribution of columns containing gaps or inconsistent residue-embedding directions.

At each guide-tree merge, the two child profiles are globally aligned using affine-gap NW with user-configurable affine gap penalties (default gap-open and gap-extension penalties of −0.5 and 0, respectively). The traceback path is applied to every sequence in the child profiles by inserting the corresponding gap characters. The aligned child profiles are then combined into a parent profile according to:

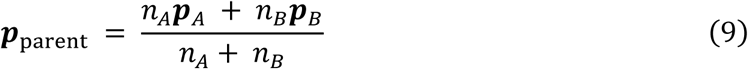

where *n_A_* and *n_B_* are the numbers of sequences represented by the two child profiles and gaps contribute zero vectors. Profile merging continues until all sequences are incorporated into a single alignment, which is written in FASTA format.

When the input network does not contain all *N*(*N* − 1)/2 interactions, missing distances are estimated using the length-adjusted cosine similarities of pooled per-residue embeddings defined in Equation 6. An isotonic regression model is fitted between estimated similarities and the scores of the computed pairs. The fitted model is then used to predict similarity scores for missing pairs, while the pre-computed scores are retained for computed pairs. The completed distance matrix is used to construct the alignment guide tree.

### Example Workflow Procedure

A fold-type IV PLP-dependent enzyme sequence set containing 12917 sequences was obtained from the oTAED database [16]. The sequence set was combined with additional sequences from the RCSB PDB database that have solved structures serving as MSA structural references [17]. Upon deduplication, a final sequence set containing 12927 sequences was obtained.

The final sequence set was passed to *Sanitize_Sequences.py* to replace unsupported characters in sequences and sequence headers. The sanitized sequence set was then used as input to *Gen-erate_Embeddings.py* and *Align_Substitution_Matrix.py* to obtain per-residue embeddings and all-vs-all negative log-transformed E-values, respectively. The ESMC 6B embeddings were obtained through the BioHub API, whereas E1 embeddings were generated locally in a separate Linux environment in retrieval-augmented mode (described in detail below). The per-residue embeddings were saved in 16-bit floating-point precision and passed to *Align_Similarity_Ma-trix.py* for all-vs-all similarity score calculations using local and global gap penalties of −2.0 and 0, respectively. The E1 embedding generation used ESMC 6B similarity scores as an input. For each sequence, 50 homologs were identified by ranking the ESMC 6B similarity scores against the target sequence and selecting sequences from the highest down with a stride of 20 (i.e., sequences with 20^th^, 40^th^, 60^th^, …, 1000^th^ highest similarity scores from the target sequence were selected as homologs). The MUSCLE super5, CLUSTALO and MAFFT MSAs were computed directly on the sanitized sequence set. Embedding-based MSAs were generated using the per-residue embeddings and corresponding similarity scores as inputs. The guide trees were built on alignment-length-normalized global similarity scores using UPGMA with 100 perturbed replicates at normalized noise scale of 0.02. An affine gap penalty of −0.5 open and 0 extend was applied. The final SSN layouts were generated based on negative log-transformed E-values and alignment-length-normalized global similarity scores using default physics and convergence parameters.

### Embedding-based MSA Benchmarking

The reference MSA was generated from 27 fold-type IV PLP-dependent enzymes with solved structures using the structural alignment (Expresso) in the T-COFFEE suite [32]. The full MSAs containing 12927 sequences were stripped to contain only the reference set with gap-only columns removed. Each stripped MSA was compared to the reference, and SP F1 and TC scores of each comparison were measured.

### AI Disclosure Statement

Generative AI tools, ChatGPT (OpenAI) and Gemini (Google), were used during development of the software to assist with code generation, debugging, refactoring, feature implementation, and development of portions of the computational workflow. Suggestions generated by the generative AI were critically evaluated by the authors, and all incorporated code and analytical procedures were subsequently reviewed and tested. Generative AI did not independently determine the study design, evaluate the resulting EMAP-SSN platform, or formulate the scientific interpretations of this work. The authors take full responsibility for the final software, computational analyses, and reported results.

## Acknowledgements

We would like to thank Owen Mototsune (CNRS) and Miguel Tsai (University of Toronto) for software testing. This work was funded by the NSERC CREATE for BioZone project (Grant# 528163) and the NSERC Alliance (ALLRP 602210-24)

## Data Availability Statement

The data that support the example workflow in this study are openly available on Zenodo at 10.5281/zenodo.23420957. The fold-type IV PLP-dependent enzyme sequence set was derived from protein sequence data obtained from the oTAED and RCSB PDB databases, as described in the Methods. The Zenodo repository contains the processed sequence dataset, per-residue protein embeddings, all-vs-all sequence similarity scores and negative log-transformed E-values, MSAs, SSN layout cache files, and other files required to reproduce the analyses presented in this study. The source code for EMAP-SSN is publicly available at https://github.com/Xuebin-Feng/EMAP-SSN, and the version used in this study is archived at 10.5281/zenodo.23417138.

## References

[1] H. J. Atkinson, J. H. Morris, T. E. Ferrin, and P. C. Babbitt, “Using sequence similarity networks for visualization of relationships across diverse protein superfamilies,” PLoS One, vol. 4, no. 2, 2009, doi: 10.1371/journal.pone.0004345.

[2] J. A. Gerlt et al., “Enzyme function initiative-enzyme similarity tool (EFI-EST): A web tool for generating protein sequence similarity networks,” Aug. 01, 2015, Elsevier B.V. doi: 10.1016/j.bbapap.2015.04.015.

[3] L. McInnes, J. Healy, and J. Melville, “UMAP: Uniform Manifold Approximation and Projection for Dimension Reduction,” Sep. 2020, [Online]. Available: http://arxiv.org/abs/1802.03426

[4] C. Camacho et al., “BLAST+: Architecture and applications,” BMC Bioinformatics, vol. 10, Dec. 2009, doi: 10.1186/1471-2105-10-421.

[5] S. F. Altschul et al., “Gapped BLAST and PSI-BLAST: a new generation of protein database search programs,” Oxford University Press, 1997. [Online]. Available: https://aca-demic.oup.com/nar/article/25/17/3389/1061651

[6] B. Rost, “Twilight zone of protein sequence alignments,” 1999. [Online]. Available: https://academic.oup.com/peds/article/12/2/85/1550637

[7] K. Kaminski, J. Ludwiczak, K. Pawlicki, V. Alva, and S. Dunin-Horkawicz, “pLM-BLAST: distant homology detection based on direct comparison of sequence representations from protein language models,” Bioinformatics, vol. 39, no. 10, Oct. 2023, doi: 10.1093/bioinformatics/btad579.

[8] S. Dash, S. R. Rahman, H. M. Hines, and W. C. Feng, “iBLAST: Incremental BLAST of new sequences via automated e-value correction,” PLoS One, vol. 16, no. 4 April 2021, Apr. 2021, doi: 10.1371/journal.pone.0249410.

[9] Z. Lin et al., “Evolutionary-scale prediction of atomic-level protein structure with a language model,” 2023. doi: 10.1126/science.ade2574.

[10] A. Rives et al., “Biological structure and function emerge from scaling unsupervised learning to 250 million protein sequences”, doi: 10.1073/pnas.2016239118/-/DCSupple-mental.

[11] C. Dallago et al., “Learned Embeddings from Deep Learning to Visualize and Predict Protein Sets,” Curr. Protoc., vol. 1, no. 5, May 2021, doi: 10.1002/cpz1.113.

[12] L. Pantolini, G. Studer, J. Pereira, J. Durairaj, G. Tauriello, and T. Schwede, “Embedding-based alignment: combining protein language models with dynamic programming alignment to detect structural similarities in the twilight-zone,” Bioinformatics, vol. 40, no. 1, Jan. 2024, doi: 10.1093/bioinformatics/btad786.

[13] R. C. Edgar, “Muscle5: High-accuracy alignment ensembles enable unbiased assessments of sequence homology and phylogeny,” Nat. Commun., vol. 13, no. 1, Dec. 2022, doi: 10.1038/s41467-022-34630-w.

[14] F. Steffen-Munsberg et al., “Bioinformatic analysis of a PLP-dependent enzyme superfamily suitable for biocatalytic applications,” Sep. 01, 2015, Elsevier Inc. doi: 10.1016/j.biotechadv.2014.12.012.

[15] M. Höhne, S. Schätzle, H. Jochens, K. Robins, and U. T. Bornscheuer, “Rational assignment of key motifs for function guides in silico enzyme identification,” Nat. Chem. Biol., vol. 6, no. 11, pp. 807–813, 2010, doi: 10.1038/nchembio.447.

[16] O. Buß, P. C. F. Buchholz, M. Gräff, P. Klausmann, J. Rudat, and J. Pleiss, “The ω-transaminase engineering database (oTAED): A navigation tool in protein sequence and structure space,” *Proteins: Structure*, Function and Bioinformatics, vol. 86, no. 5, pp. 566– 580, May 2018, doi: 10.1002/prot.25477.

[17] H. M. Berman et al., “The Protein Data Bank,” 2000. [Online]. Available: http://www.rcsb.org/pdb/status.html

[18] S. Candido et al., “Language Modeling Materializes a World Model of Protein Biology,” Jun. 04, 2026. doi: 10.64898/2026.06.03.729735.

[19] S. Jain, J. Beazer, J. A. Ruffolo, A. Bhatnagar, and A. Madani, “E1: Retrieval-Augmented Protein Encoder Models,” Nov. 13, 2025. doi: 10.1101/2025.11.12.688125.

[20] V. A. Traag, L. Waltman, and N. J. van Eck, “From Louvain to Leiden: guaranteeing well-connected communities,” Sci. Rep., vol. 9, no. 1, Dec. 2019, doi: 10.1038/s41598-019-41695-z.

[21] A. S. Rose, G. Tomasello, Á. S. Kovács, L. Autin, and D. Sehnal, “Mol* web molecular graphics engine,” Protein Science, vol. 35, no. 4, Apr. 2026, doi: 10.1002/pro.70514.

[22] M. Blum et al., “InterPro: The protein sequence classification resource in 2025,” Nucleic Acids Res., vol. 53, no. D1, pp. D444–D456, Jan. 2025, doi: 10.1093/nar/gkae1082.

[23] A. Kochevenko, H. J. Klee, A. R. Fernie, and W. L. Araújo, “Molecular identification of a further branched-chain aminotransferase 7 (BCAT7) in tomato plants,” J. Plant Physiol., vol. 169, no. 5, pp. 437–443, Mar. 2012, doi: 10.1016/j.jplph.2011.12.002.

[24] T. Meinnel, C. Lazennec, S. Villoing, and S. Blanquet, “Structure-Function Relationships within the Peptide Deformylase Family. Evidence for a Conserved Architecture of the Active Site Involving Three Conserved Motifs and a Metal Ion,” 1997.

[25] S. A. Shilova et al., “To the Understanding of Catalysis by D-Amino Acid Transaminases: A Case Study of the Enzyme from Aminobacterium colombiense,” Molecules, vol. 28, no. 5, Mar. 2023, doi: 10.3390/molecules28052109.

[26] M. Voss et al., “Creation of (R)-Amine Transaminase Activity within an α-Amino Acid Transaminase Scaffold,” ACS Chem. Biol., vol. 15, no. 2, pp. 416–424, Feb. 2020, doi: 10.1021/acschembio.9b00888.

[27] D. Peisach, D. M. Chipman, P. W. Van Ophem, J. M. Manning, and D. Ringe, “Crystallographic Study of Steps along the Reaction Pathway of D-Amino Acid Aminotransferase.” doi: 10.1021/bi972884d.

[28] F. Sievers et al., “Fast, scalable generation of high-quality protein multiple sequence alignments using Clustal Omega,” Mol. Syst. Biol., p. 539, 2011, doi: 10.1038/msb.2011.75.

[29] K. Katoh and D. M. Standley, “MAFFT multiple sequence alignment software version 7: Improvements in performance and usability,” Mol. Biol. Evol., vol. 30, no. 4, pp. 772–780, Apr. 2013, doi: 10.1093/molbev/mst010.

[30] S. Mirarab and T. Warnow, “FASTSP: Linear time calculation of alignment accuracy,” Bioinformatics, vol. 27, no. 23, pp. 3250–3258, Dec. 2011, doi: 10.1093/bioinformatics/btr553.

[31] J. D. Thompson, F. Plewniak, and O. Poch, “BAliBASE: a benchmark alignment database for the evaluation of multiple alignment programs.” [Online]. Available: http://www-ig-bmc.u-strasbg.fr/BioInfo/

[32] F. Armougom et al., “Expresso: Automatic incorporation of structural information in multiple sequence alignments using 3D-Coffee,” Nucleic Acids Res., vol. 34, no. WEB. SERV. ISS., 2006, doi: 10.1093/nar/gkl092.

[33] M. Heinzinger et al., “Bilingual language model for protein sequence and structure,” NAR Genom. Bioinform., vol. 6, no. 4, Dec. 2024, doi: 10.1093/nargab/lqae150.

[34] N. Brandes, D. Ofer, Y. Peleg, N. Rappoport, and M. Linial, “ProteinBERT: a universal deeplearning model of protein sequence and function,” Bioinformatics, vol. 38, no. 8, pp. 2102–2110, Apr. 2022, doi: 10.1093/bioinformatics/btac020.

[35] A. Elnaggar et al., “Ankh ☥: Optimized Protein Language Model Unlocks General-Purpose Modelling,” Jan. 18, 2023. doi: 10.1101/2023.01.16.524265.

