## supplementary material for "EMAP-SSN: An Embedding- and Multiple-Alignment-Integrated Sequence Similarity Network Platform for Interactive Exploration of Protein Sequence Space"

### Supplementary Figures and Tables

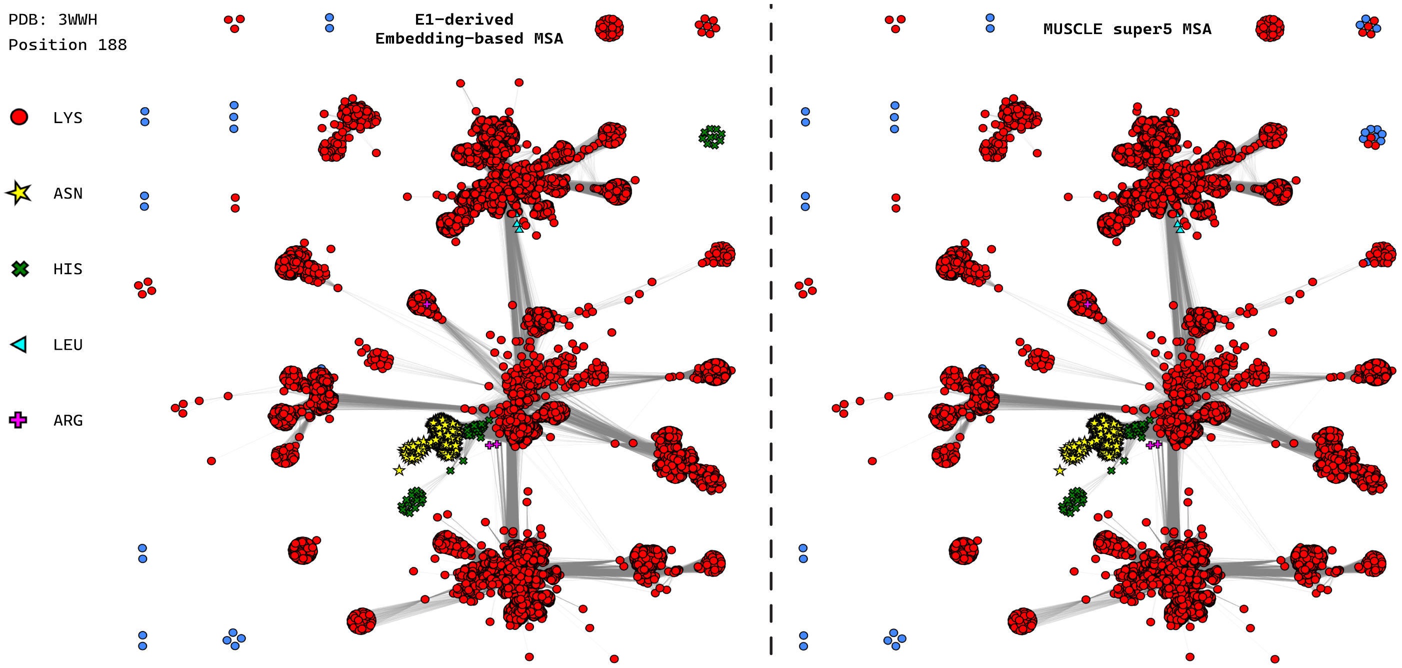


Figure S1. E1-based SSN loaded with E1-derived embedding-based MSA and MUSCLE super5 MSA colored based on the amino acid identity at the conserved lysine position (188 in the 3WWH numbering scheme).

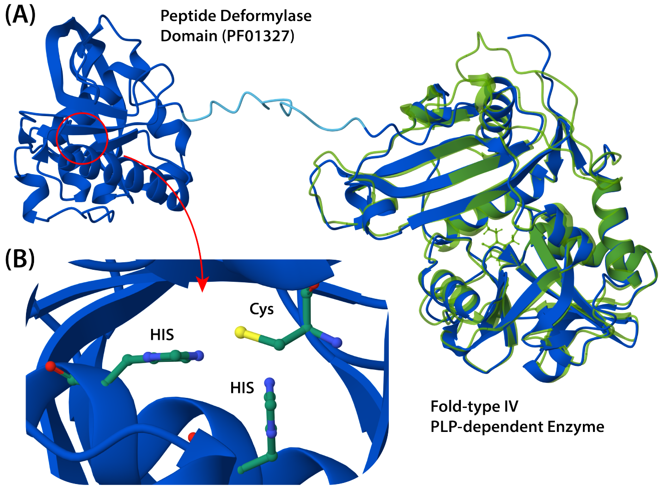


Figure S2. (A) ESM3-predicted (98B via BioHub API) structure of a fold-type IV PLP-dependent enzyme from *Burkholderia* sp. with a long N-terminal extension (blue) superimposed on 3WWH (green). (B) Active site residues of the peptide deformylase domain.

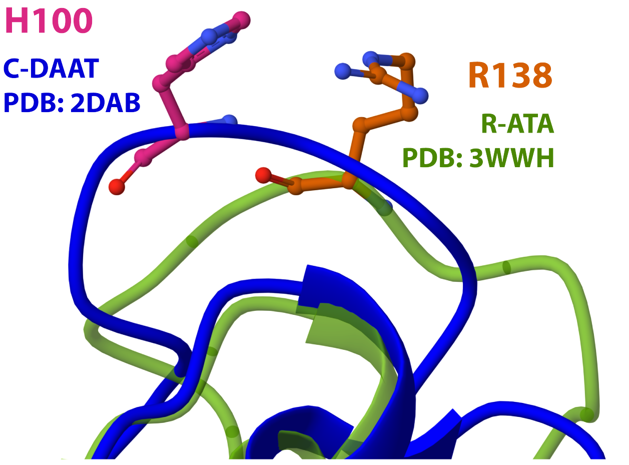


Figure S3. Structural alignment of H100 of 2DAB (C-DAAT) and R138 of 3WWH (R-ATA). The two positions are on flexible loops and cannot be aligned cleanly. The same applies to R98 of 2DAB and P135 of 3WWH (comparison not shown.

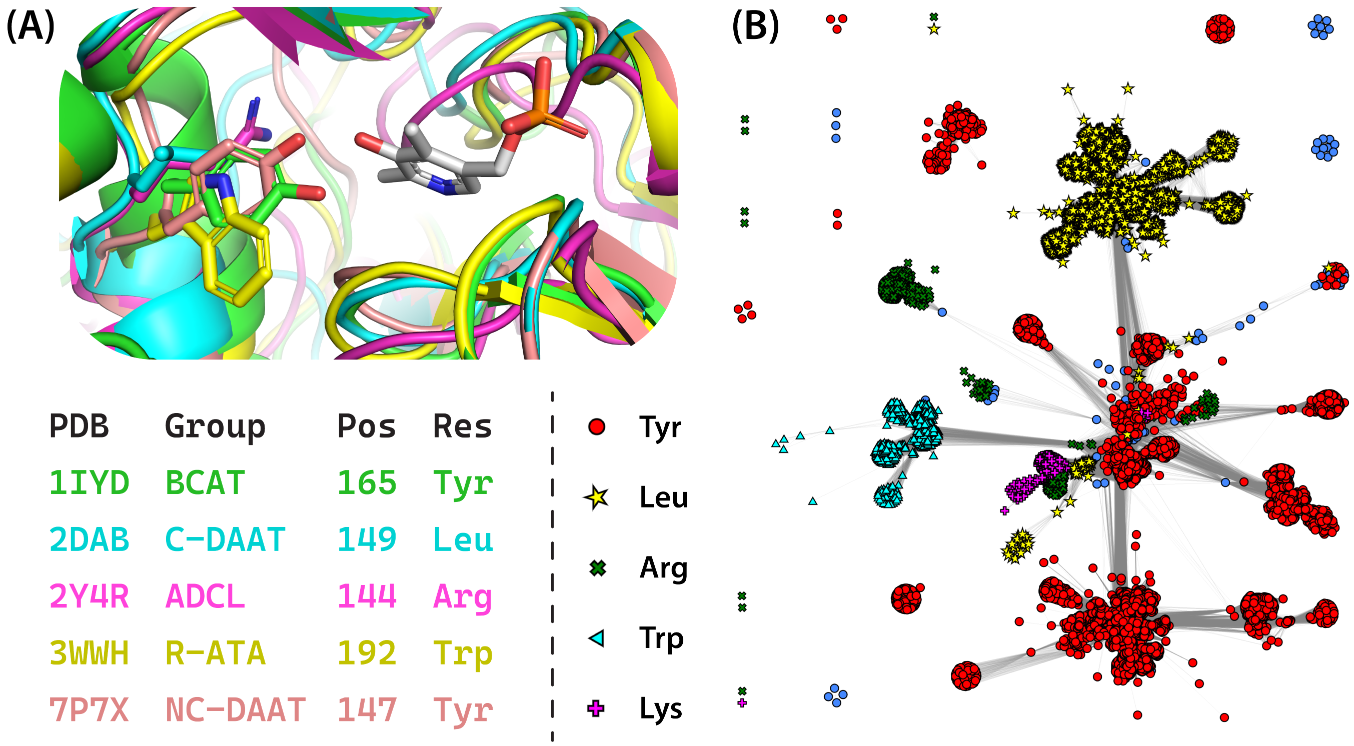


Figure S4. (A) Alignments of five PDB structures each from a distinct functional group with amino acids at the 3WWH position 192 highlighted. Although not included in the active site signature region, position 192 points directly toward the 3'-hydroxyl of the PLP cofactor. (B) The E1-derived SSN colored based on the amino acid identity at the 3WWH position 192. Different amino acid identities showed clear cluster-based separation.

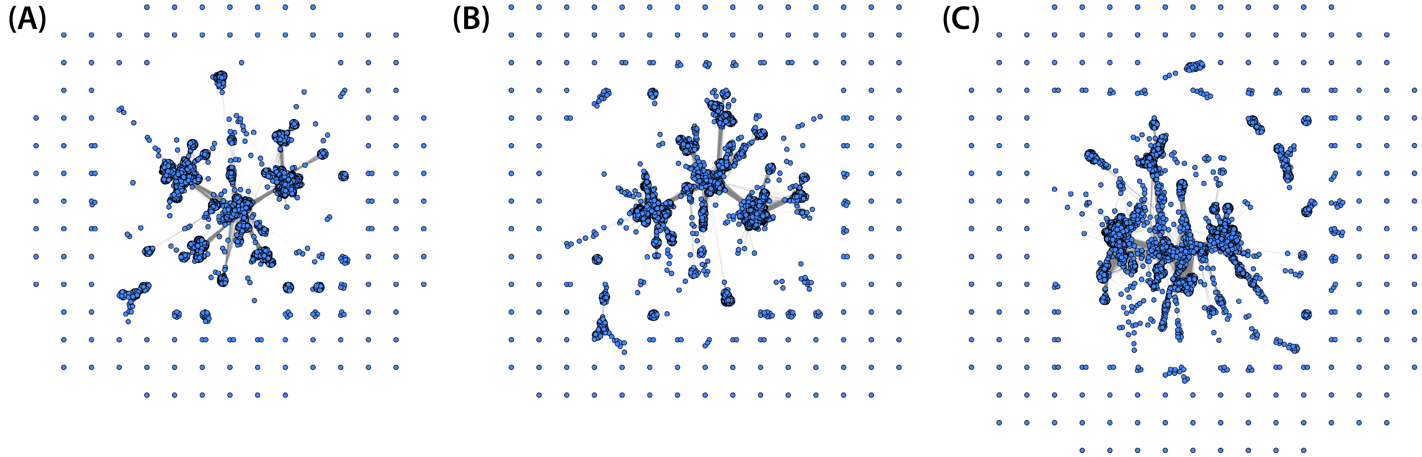


Figure S5. Initial layouts of E1-based SSNs generated with similarity networks with different sparsity levels. (A) 25% sparse, the network retains a similar topology to the SSN generated with the full network. (B) 50% sparse, the network begins to fractionate, forming more free nodes and small clusters. (C) 75% sparse, the network still retains the overall topology; however, individual clusters begin to lose their shapes and form long chains of nodes.

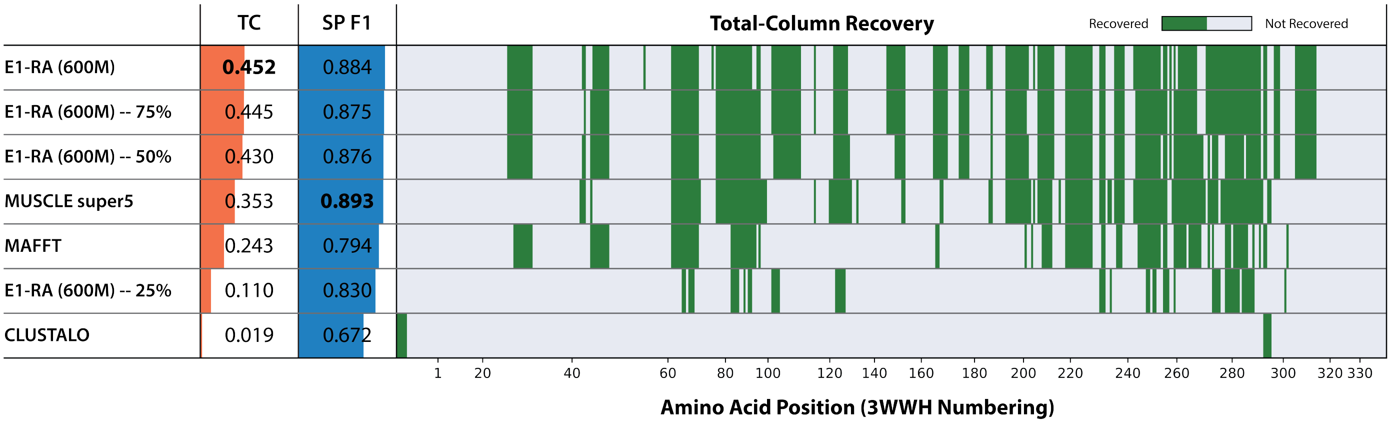


Figure S6. Total column (TC) and sum of pair F1 (SP F1) scores of alignments showing the effect of network sparsity on the accuracy of the embedding-based MSA calculation. The accuracy was not affected much until the sparsity increased above 50% (or computed pairs / total pairs drops below 50%), at which point the accuracy drops dramatically.

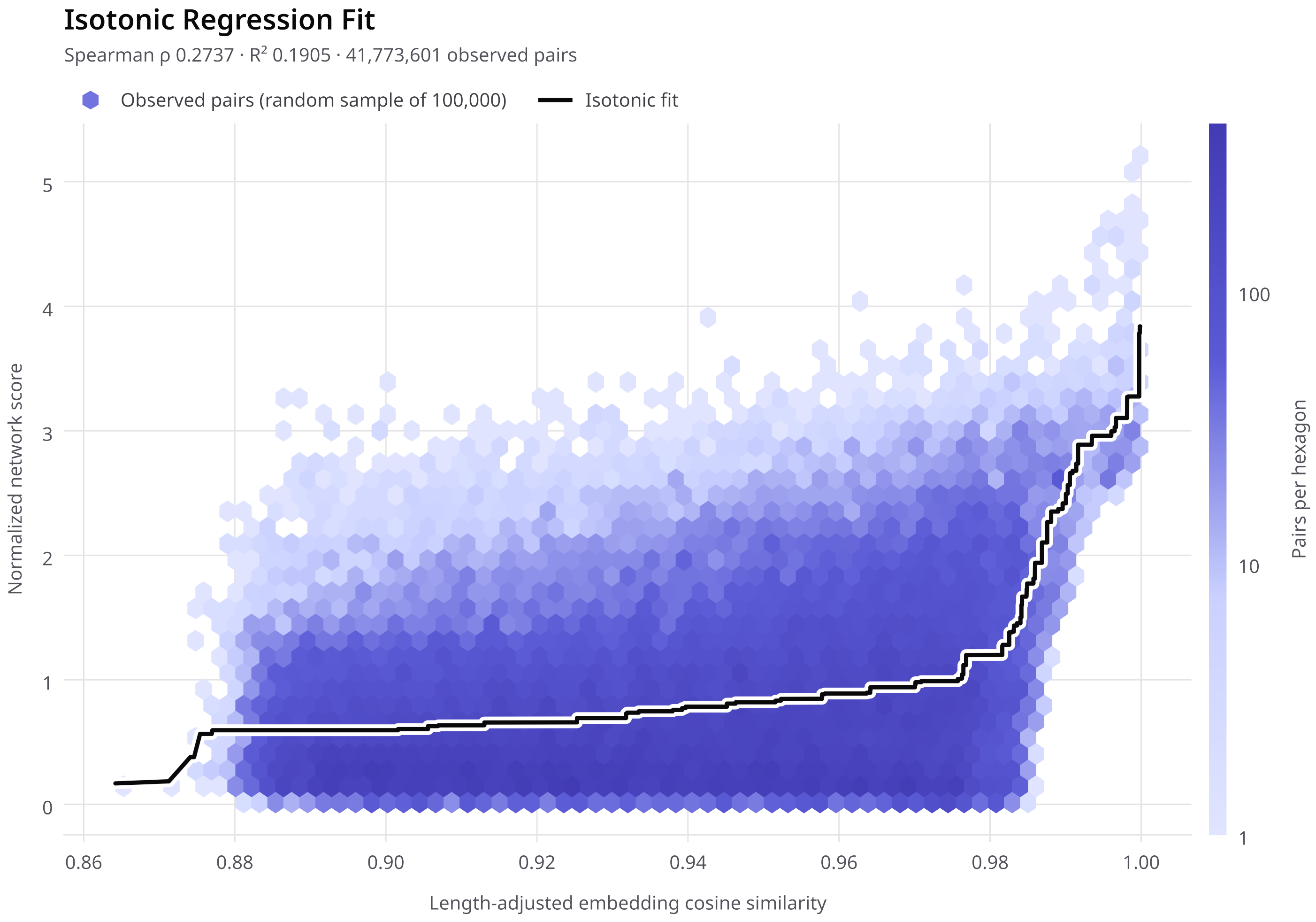


Figure S7. Isotonic regression plotted between alignment-length-normalized global alignment score and the max-pooled and distance corrected cosine similarities of the same sequence pairs. The correlation is more significant on the higher end of the length adjusted cosine similarities.

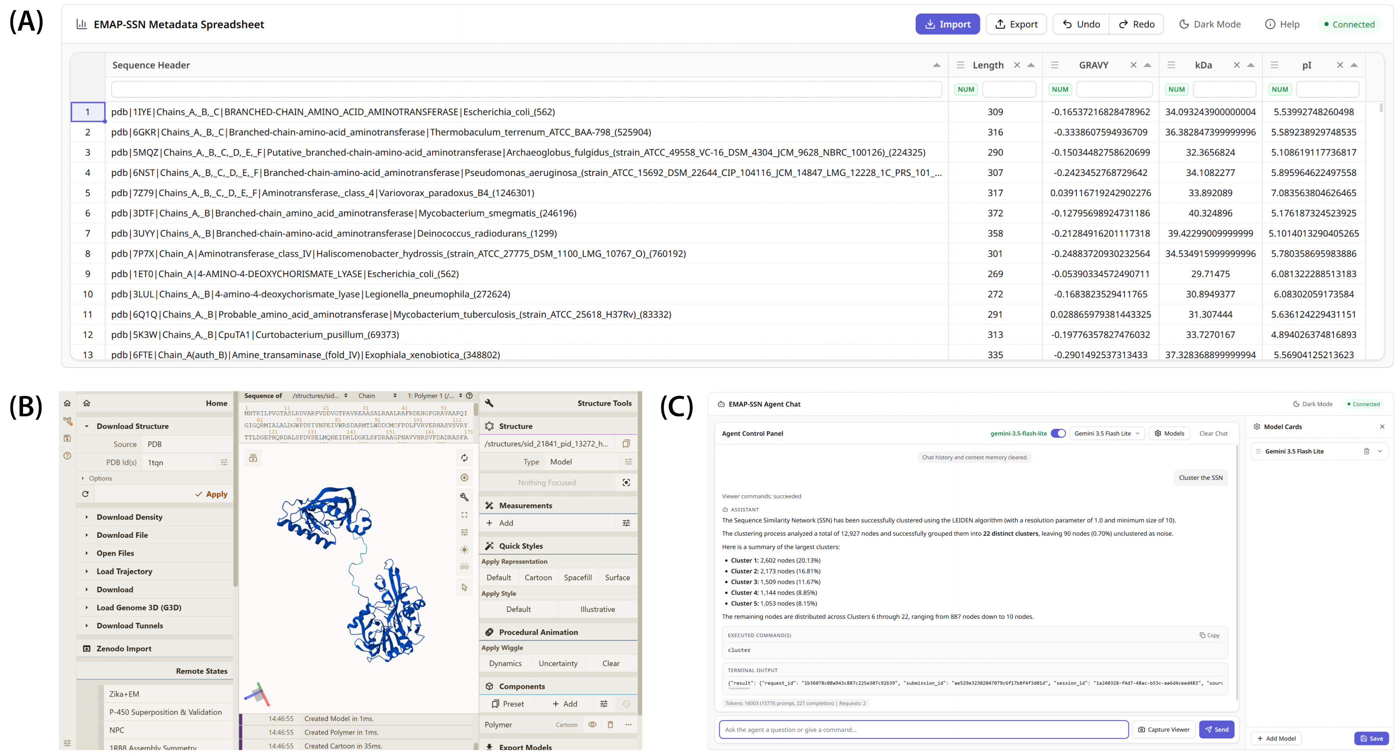


Figure S8. The three web-based tools hosted by the local HTTP server. (A) The Tabulator-based metadata viewer. (B) The integrated Mol* web interface for viewing structures generated by the ESMFOLD command. (C) The agent chat web interface allowing users to connect external APIs to allow chatbots to translate natural-language requests into CLI commands.

### Corresponding Sequences of Structures shown in Figure 4 and Figure S2

#### Figure 4 – Protein containing Sulfotransferase 5 Domain N-terminal Extension

```
>sid|119541|pid|69403|hfid|11|sfid|2|gb|KHN16146.1|gi|734362462|taxonID|3848|Glycinesoja

MEVIHLWSAPRSLSTTVMYSFAQRDDIEVLDEPLYANFLRVTAVHRPYKEELLSKMESDGNKVVKDIIYRPGNSKYRFCKHMSKQRILGLPE
DLMKKGKHFILIRNPLDILPSFDEVVPPSFFELGLAELVCIYNELCEIGKPPPVVDAAELQQDPEATLRALCNDLEIPFQPAMLNWEAGPKP
IDGLWAPWWYKTVHKSTGFKEENKYPQPFPFSLYNLLEQSLPLYNMLRRHVKKKPSLLGTPLPTPDLPVPANEKLLAWVGNEIVPRESAKVS
VFDSVVQGGDSVWEGLRVYNGKIFKLEEHLDRMFDSAKALAFENVPTRDKIKEAIFKTLIRNGMFDNSHIRLSLTRGKKVTSGMSPTLNLYG
CTLIVLAEWKPPVYDNEHGIVLVTATTRRNSPNNLDSKIHHNNLLNNILAKIEGNNAKADDAIMLDKDGYVSETNATNMFIVKRGAVLTPHA
DYCLPGITRATVMDLVVKEQFILEERRISLSEVHTADEVVKVDGRIIGNGKVGPVTRQLQAAYRKLTEQLGVPISNYLEA
```

#### Figure S2 – Protein containing Peptide Deformylase Domain N-terminal Extension

```
>sid|21841|pid|13272|hfid|10|sfid|2|gi|915395138|taxonID|95486|Burkholderiacenocepacia|
gb|ACA94085.1|gi|169819503|taxonID|406425|BurkholderiacenocepaciaMC0-3

MNTRILPVGTASLRDVARPVDDVGTPAVREAASALRAALRAFRDEHGFGRAVAAPQIGIGQRMIALALDGWPDVIVNPEIVWRSDARMTLWD
DCMCFPDLFVRVERHASVSVRYTTLDGEPHQRDALSPDVSELMQHEIDHLDGKLSFDRAAGPNAVVHRSVFDADRASFAAQVDYTPNVPRET
RMAHRGIPSVEAPGFPQGAAYMNGRFIPIADARVSVLDWGFLHSDVTYDTVHVWNGRFFRLDKHIERFRRSLARLRLNVPLTDDALRDILVE
CVRRSGLRHAYVEMLCTRGVSPTFSRDPRDAVNQFIAFAVPYGSVANERQLREGLHLHVIDDVRRIPPESVDPQIKNYHWLDLVAGLLKGYD
AGAESVLLKCTDGSIAEGPGFNVFVVRDGRLRTPERGVLHGITRQTVFELATAMGIDAQAARIDDAQLRDADEVFITSTAGGIMPVTRLNDA
TIGDGRPGPVTRRLFDAYWAKHGDPAWSLAVDYADG
```

### CLI Commands used to Generate Figures and Data

Figure 2 – Initial Layouts

First, maximize the "Viewer" GUI or enter full-screen mode, and adjust view to a fixed zoom level by `zoom 200`

Then print the entire plotting area by `print full transparent`

Figure 3 – Clustering and Conserved Position Coloring

First hide free/isolated nodes by `hide free`

Automatic clustering can be done by `cluster 1.0 20`

Then print the entire plotting area by `zoom 100` and then `print full transparent`

For coloring, first reset SSN to start fresh `reset color`

Switch the reference to 3WWH `reference 3WWH`

use a single COLOR command to specify color, shape and size of each node

```
color K188 red N188 yellow * x1.5 H188 green x L188 cyan ^ R188 magenta +
```

Then print the entire plotting area by `zoom 100` and then `print full transparent`

Figure 4 – Color SSN based on Sequence Length and then Predict Protein Structure

First reset SSN and node order, which was changed by the COLOR command `reset color shape size order`

Color the SSN based on sequence length `spectrum {length}`

Then manually zoom into the area shown in the figure and print the viewing area `print transparent`

To generate the predicted fold, first manually select the node to predict by right-clicking it. Then use `esmfold large`

The calculation is routed through BioHub API. To run locally, use `esmfold` without an argument.

Figure 6 – Sequence Logo and Cluster 3 Coloring

The panel A logo generation can be done using a Python script interpreted by the RUN command.

First create the custom groups Active-ASN and Active-HIS by manually selecting the nodes and calling `group <group_name>`

Then create a Python file with the following content

```
# Loop settings
positions = "[58,62,67,69,71,122,124,126,135,138,189,192,194,223,242,282]"
func_groups = ["BCAT", "C-DAAT", "NC-DAAT", "R-ATA", "ADCL", "Active-ASN", "Active-HIS"]
print(f"logo {positions} 0.9 Global_logo.svg")
# Loop execution
for group in func_groups:
    print(f"logo #{group}# {positions} 0.9 {group}_logo.svg")
```

Then use the RUN command by simply calling `run` and select the file.

For panel B, first hide nodes which are not a part of the four clusters

`hide !(#cluster_2#|#cluster_3#|#cluster_14#|#cluster_18#)`

Then take a zoomed-in picture `zoom 60` and `print full transparent`

For panel C, only show nodes in cluster 3 and apply additional colors `hide !#cluster_3#`

`color F62&F138 red M62&F138 yellow L62&H138 green`
